# Population coding of time-varying sounds in the non-lemniscal Inferior Colliculus

**DOI:** 10.1101/2023.08.14.553263

**Authors:** Kaiwen Shi, Gunnar L. Quass, Meike M. Rogalla, Alexander N. Ford, Jordyn E. Czarny, Pierre F. Apostolides

**Affiliations:** Kresge Hearing Research Institute, Department of Otolaryngology — Head & Neck Surgery, University of Michigan Medical School, Ann Arbor, MI, 48109; Department of Molecular & Integrative Physiology, University of Michigan Medical School, Ann Arbor, MI, 48109

## Abstract

The inferior colliculus (IC) of the midbrain is important for complex sound processing, such as discriminating conspecific vocalizations and human speech. The IC’s non-lemniscal, dorsal “shell” region is likely important for this process, as neurons in these layers project to higher-order thalamic nuclei that subsequently funnel acoustic signals to the amygdala and non-primary auditory cortices; forebrain circuits important for vocalization coding in a variety of mammals, including humans. However, the extent to which shell IC neurons transmit acoustic features necessary to discern vocalizations is less clear, owing to the technical difficulty of recording from neurons in the IC’s superficial layers via traditional approaches. Here we use 2-photon Ca^2+^ imaging in mice of either sex to test how shell IC neuron populations encode the rate and depth of amplitude modulation, important sound cues for speech perception. Most shell IC neurons were broadly tuned, with a low neurometric discrimination of amplitude modulation rate; only a subset were highly selective to specific modulation rates. Nevertheless, neural network classifier trained on fluorescence data from shell IC neuron populations accurately classified amplitude modulation rate, and decoding accuracy was only marginally reduced when highly tuned neurons were omitted from training data. Rather, classifier accuracy increased monotonically with the modulation depth of the training data, such that classifiers trained on full-depth modulated sounds had median decoding errors of ∼0.2 octaves. Thus, shell IC neurons may transmit time-varying signals via a population code, with perhaps limited reliance on the discriminative capacity of any individual neuron.

**Significance Statement:** The IC’s shell layers originate a “non-lemniscal” pathway whose first- and second-order targets are thought important for perceiving conspecific vocalizations and human speech. However, prior studies suggest that individual shell IC neurons are broadly tuned and have high response thresholds, implying a limited reliability of efferent signals. Here we use Ca^2+^ imaging to test how shell IC neurons encode amplitude modulation, a key sound cue for speech perception and stream segregation. We show that the rate and depth of amplitude modulation is accurately represented in the ensemble activity of shell IC neuron populations. Thus, downstream targets can read out a sound’s temporal envelope from a distributed rate code transmitted by populations of broadly tuned neurons.

## Introduction

The “amplitude modulation” (AM) of a sound’s envelope is a hallmark of vocal communication across species, and supports important cognitive processes such as stream segregation and intelligibility of syllabic features (Drullman et al., 1994; Shannon et al., 1995; Zeng et al., 2005; Gallun and Souza, 2008; Elliott and Theunissen, 2009; Koumura et al., 2019, 2023). Despite this importance, how the brain reliably encodes AM sounds, and thus may provide a key building block of receptive language, remains poorly understood.

The inferior colliculus (IC) is an evolutionarily conserved midbrain structure that plays a pivotal role in AM perception (Champoux et al., 2007), and the first major central auditory structure where neuronal responses to AM sounds change from a temporal- to a rate-based representation (Hewitt and Meddis, 1994; Lorenzi et al, 1995; Krishna and Semple, 2000; Nelson and Carney, 2007; Tan and Borst, 2007; Ter-Mikaelian et al., 2007; Dicke et al., 2007; Geis and Borst, 2009; Wang et al., 2022). Indeed, although neurons in sub-tectal brainstem nuclei phase-lock their firing rates to AM fluctuations, mean spike rates are often insensitive to the temporal characteristics of the sound envelope (Frisina et al., 1990; Rhode and Greenberg, 1994; Zhao and Liang, 1994). By contrast, mean spike rates of IC neurons consistently and dramatically vary depending on a sound’s AM pattern, with specific IC neurons often displaying sharp AM rate selectivity (Langner and Schreiner, 1988; Krishna and Semple, 2000; Nelson and Carney, 2007). Compared with a temporal code which requires sampling over multiple period cycles, such rate coding across spatially distributed neurons might facilitate more efficient representation of sound features (Bendor and Wang, 2007; Wang et al., 2008) and drive behavior (Bagur et al., 2023). The IC is also the first bifurcation site of primary and “higher-order” acoustic pathways. Neurons in the IC’s central nucleus project to the primary auditory thalamus, the ventral medial geniculate body (MGB). By contrast, neurons located in the surrounding dorsomedial and lateral “shell” IC nuclei project to the higher-order dorsal and medial MGB (Oliver and Morest, 1984; Winer et al., 1998; Mellott et al., 2014; Xiong et al., 2015; Chen et al., 2018; Cai et al., 2019; Oberle et al., 2022). This divergence is notable, as higher-order MGB nuclei relay acoustic signals to the amygdala, non-primary auditory cortices, and the striatum (LeDoux et al., 1991; Bordi and LeDoux, 1994; Lee and Winer, 2008; Ponvert and Jaramillo, 2019). These forebrain circuits are important for processing conspecific vocalizations and human speech (Gadziola et al., 2012; Grimsley et al., 2013; Gadziola et al., 2016; Hamilton et al., 2021), as well as orchestrating behavioral responses to ethologically relevant sounds (Miura et al., 2020; Carcea et al., 2021; Valtcheva et al., 2023).

Despite prominent connections with behaviorally relevant circuits, little is known about how shell IC neurons respond to AM sounds; most studies of AM coding have been conducted in the central IC. This knowledge gap may be due to shell IC neurons’ high response thresholds, loose tonotopy, and variable response fidelity compared to central IC neurons (Syka et al., 2000; Lumani and Zhang, 2010; Barnstedt et al., 2015; Parras et al., 2017; Wong and Borst, 2019; Chen and Song, 2019). Additionally, shell IC neurons’ precarious location at the superficial tectal layers complicates the use of single neuron recording techniques to study a variably active neuronal population. Rather, a tacit assumption is that non-lemniscal auditory system eschews high-fidelity representations of acoustic features in favor of coarser signals potentially related to a sound’s contextual novelty (Ayala et al., 2015; Parras et al., 2017) or behavioral salience (Olds et al., 1972; Edeline and Weinberger, 1992; Poremba and Gabriel, 2001; Leppla et al., 2023). Here we use 2-photon Ca^2+^ imaging to study how mouse shell IC neurons respond to sinusoidally amplitude modulated (sAM) sounds. We find that all the major sAM tuning properties previously described in the central IC – low-pass, high-pass, band-pass, and band-reject – are also found in the shell IC. Increasing the sAM depth monotonically enhanced activity, indicating limited mixed selectivity to specific combinations of sAM rate and -depth. Although individual shell IC neurons often displayed low neurometric selectivity for sAM sounds, decoder algorithms trained on shell IC fluorescence data could accurately classify sAM features with high reliability. Thus, a population code enables the IC’s non-lemniscal efferents to can transmit detailed information regarding time-varying acoustic features.

## Materials & Methods

### Surgeries

Experiments were approved by Michigan Medicine’s IACUC and performed in accordance with NIH’s guide for the care and use of laboratory animals. Surgical procedures were performed under aseptic conditions with 5-8-week-old male and female C57/Bl6J mice ordered from Jackson Labs (N = 9, 4 female and 5 male) or offspring of VGAT-ires-cre × Ai14 breeders in our colony (N = 4, 2 female and 2 male). Mice were anesthetized with 4-5% isoflurane delivered in O^2^ and mounted in a stereotaxic apparatus (David Kopf Instruments) and injected with carprofen as pre-surgical analgesic. Body temperature was maintained near 37 C° with a feedback-controlled homeothermic blanket (Harvard Apparatus). Isoflurane concentration was subsequently reduced to 1.5-3 %, the scalp was cleaned with iodine solution (betadine), shaved, and subsequently removed. The periosteum was carefully removed and the exposed skull was etched with a sharp scalpel. A ∼2 mm circular craniotomy was carefully drilled over the left IC using a Foredom microdrill (0.9 mm caudal and 1.0 mm lateral from the lambdoid sutures), employing multiple rounds of short (<10 s) drilling, interleaved with ∼1-2 min application periods of chilled phosphate buffered saline (PBS) to the skull. Care was taken so as to not damage the transverse sinus bordering the IC and superior colliculus during the surgery. Following removal of the skull, 12.5-25 nL of virus solution was injected at 3-6 sites in the IC (∼100 nL total volume). A custom-made cranial window comprised of three stacked 2 mm diameter coverslips adhered to a 4 mm diameter coverslip (Potomac Glass) was inserted into the craniotomy and held in place with gentle pressure. The cranial window was bonded to the skull with cyanoacrylate glue (Loctite) and dental cement (Ortho-Jet). The skull was then covered in a thin layer of glue and a custom titanium headbar was cemented onto the skull. Mice received a post-operative injection of buprenorphine (0.03 mg/kg s.c., Par Pharmaceuticals) for analgesia and 24 hours later, a second dose of carprofen (5 mg/kg, s.c., Spring Meds). Mice were monitored for 7-10 days following surgery for any sign of discomfort.

### 2-photon Ca^2+^ imaging

Beginning 10-14 days following surgery, mice were handled by the experimenter and acclimated to increasingly prolonged durations of head-fixation while sitting quietly in a custom acrylic glass tube. For data collection, mice were head-fixed under a Janelia MIMMs style microscope (Sutter Instruments) and 2-photon excitation was delivered at 920 nm through a Nikon 16x objective focused axially through the cranial window onto the superficial layers of the left IC using a tunable laser (Coherent Chameleon). Peak laser power measured at the objective was typically 20-40 mW and did not exceed 60 mW. GCaMP and tdTomato fluorescence were collected using GaASP photomultiplier tubes (Hamamatsu). Frame scans (512 × 512 pixels, 30 Hz frame rate) were collected for 5-6 s on each trial, with a 3-5 s inter-trial interval where no laser light was delivered to the brain.

### Stimulus presentation

Acoustic stimuli were generated using a Tucker-Davis Technologies RZ6 and delivered in free field at 65-70 dB SPL via a calibrated electrostatic speaker (TDT ES1) positioned ∼10 cm from the mouse’s right ear. Stimuli with different sAM rates and depths were delivered in pseudo-random order once every 8-11 s. We typically delivered 10 repetitions of each stimulus when imaging a single FOV, which required 40-80 min to complete. 1-2 FOVs were typically imaged from a single mouse in each session, which typically required 1-2 hours of total head-fixation time per session. Subjects underwent multiple imaging sessions over the course of 2-3 weeks.

### Data analysis

Raw movies were motion-corrected using non-rigid registration, and regions of interests (ROIs) corresponding to individual cell bodies were generated to extract the raw fluorescence signal from somata and surrounding neuropil using Suite2p (Pachitariu et al., 2016). Fluorescence changes were measured by subtracting the corresponding neuropil signal (scaled between 70-100 % based on recording quality) on each trial and calculating the change in fluorescence as (F–F_0_)/F_0_ (ΔF/F). In these experiments, F_0_ represents the mean fluorescence intensity during the 0.5 - 1 s baseline period prior to sound onset on each trial.

Analyses presented in this paper are performed only on ROIs displaying a statistically significant sound response, as determined using the ‘signal autocorrelation’ bootstrapping procedure of Geis et al., (2011) and Wong and Borst (2019) and for each ROI in an FOV, the raw ΔF/F traces during, and 1 s following, sound presentation were correlated between individual repetitions of the same stimulus presentation. We calculated a mean Pearson’s ρ value across all possible pairs of trials of the same stimulus presentation, thus providing a metric of how reliably the particular sound stimulus drives a response in the neuron. We then generated “shuffled” datasets of randomly chosen sequential frames from the fluorescence time series across all trials of the same stimulus presentation. The length of these segments, and the number of pairs of trials, were always equal to the length of fluorescence traces and number of stimulus presentations used to calculate the true signal autocorrelation described above. In addition, the specific fluorescence time series was included in only a single randomly chosen segment. We calculated the signal autocorrelation for these shuffled data sets were and the process was bootstrapped 10,000 times. A p-value was then calculated for the ROI, defined as the percentage of shuffled bootstrapped iterations whose mean rho value exceeded that of the true data after applying the Bonferroni-Holm correction.

For measures of peak ΔF/F, we averaged all trials corresponding to the same sound stimulus and calculated the mean signal centered ±1 frames around the local absolute maximum of the fluorescence change during sound presentation.

### Lifetime sparseness

Lifetime sparseness was used to calculate the selectivity of single shell IC neurons to sAM rates for fully modulated and unmodulated sounds,

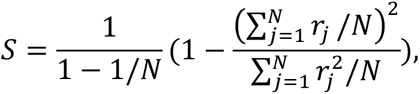

where N is the number of different sAM rates and *r_j_* is the peak of the neuronal response to sAM rate j during sound presentation. Lifetime sparseness was calculated for every individual neuron.

### Neurometric sensitivity index (d-prime)

To assess the shell IC neurons’ ability to separate different sound stimuli, we computed d-prime for all sound-responsive neurons. The neurometric d-prime for a single neuron is defined as

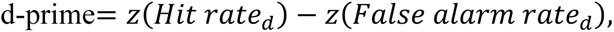

where *z* is the inverse of the normal cumulative function, and d is the sAM depth of the sound. We define a trial to be a hit if a neuron’s response at its preferred sAM rate exceeded 3 times the standard deviation during its baseline (1s before sound onset). Similarly, a trial is defined as a false alarm if that neuron’s response at non-preferred sAM rates exceeded 3 times the standard deviation during the baseline. In addition, the standard deviation σ of the difference of neurometric d-primes between 100% and 20% sAM depth for the stimuli at the preferred rate, of all neurons was calculated. If this difference was larger or smaller than 2σ or −2σ, respectively, its d-prime was defined to have either an increasing or decreasing trend.

### Convolutional neural network (CNN) decoder

A 3D-matrix containing neural population Ca^2+^ fluorescence activity from 38 sessions from 13 mice (Neurons × Frames × Trials) was passed to a CNN decoder (MATLAB_R2021b). The decoder was composed of an input layer, a convolutional layer, a max-pooling layer, a fully connected layer, and an output layer (classification or regression layer). A dropout layer and L2 regularization were added to prevent overfitting. Below we provide the details of operations of these building blocks.

#### Convolutional Layer

The convolutional layer, consisting of k kernels, performs a linear operation to extract features from the input. The following parameters mainly define the convolutional layer: *s_k_, s_p_, s_s_, k, and **W***, where *s_k_, s_p_, s_s_* are the size of the kernels, the padding, and the stride, k is the number of kernels, and ***W*** is the weight array of the kernels. The input ***X*** for the convolutional layer is a temporal representation of each neuron’s Ca^2+^ fluorescence, expressed as a 2D array (*h_in_*, *w_in_*), where the height *h_in_* is the number of neurons in each imaging session, the width *w_in_* is the evaluation time (from sound onset to 1s or 0.5s after sound offset, depending on sound duration), and each entry contains the normalized fluorescence value for a given neuron at a given time. K filtered feature maps are generated after each kernel is convolved with the input. The output of the convolutional layer 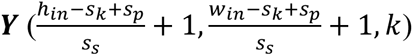 is expressed as:

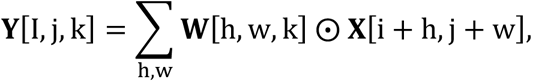

where ⊙ denotes element-wise multiplication. In our analysis, the size of the kernels is set to be 6 × 6 and the number of kernels is 32. A non-linearity is then added to the output of the convolutional layer through the rectified linear unit (ReLU) activation function, which is defined as:

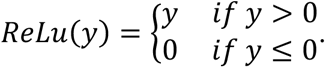

This nonlinearity enables a neural network to learn and operate with nonlinear functions. The size of the stride was 1 × 1 and we used zero-padding to avoid input/output size-discrepancies.

#### Max-Pooling Layer

The pooling layer downsamples its input via operation in a *q* × *q* region. Taking the input of size (*h_in_*, *w_in_*), a max-pooling layer returns a 2D output array 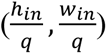 by taking the maximum value within the *q* × *q* region. Here, the size of max-pooling is 3 × 3 with a 1 × 1 stride.

#### Fully Connected (FC) Layer

The FC layer first flattens its input ***X*** (*h_in_*, *w_in_*) into a one-dimensional array (*n_out_*). This is achieved by reducing the input **X** to an array ***X_flat_***(*h_in_* × *w_in_*), and multiplying ***X****_flat_* with its weight matrix ***W***. *L*_2_regularization is incorporated in the FC layer, and the FC layer is followed by the output layer (either classification or regression).

#### Classification Layer

For classification tasks, the SoftMax layer transfers the outputs of the FC layer into a probability distribution of N labels by a normalized exponential function. Following the SoftMax layer, the classification layer adopts the cross-entropy loss function,

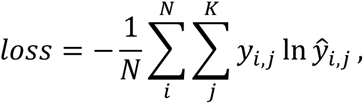

where N is the size of the training set, K is the number of classes, *ŷ* is the probability generated by the SoftMax layer and y indicates the ground truth.

#### Regression Layer

For regression tasks, the FC layer is followed by a regression layer, which adopts the mean-square-error (MSE) as a loss function,

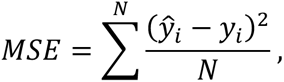

where *ŷ_i_* is the prediction generated by the network, *y_i_* is the ground truth, and N is the number of trials.

#### Dropout Layer

The dropout layer randomly sets 50 % of the weights of the FC layer to zero for one training batch, while the rest remains untouched. This operation prevents overfitting of a neural network by avoiding complicated co-adaptations of feature detectors in the training data (Hinton et al., 2012).

#### *L*_2_ Regularization

Regularization is one of the most effective ways to prevent overfitting. An *L*_2_ regularizer penalizes the loss functions by introducing a term 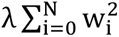 to the loss function, where w represents the weight and λ represents a coefficient determining the scale of regularization. (Cortes et al., 2012)

### Network training and evaluation

The neural network was trained to decode sAM sound parameters using neural population activity by stochastic gradient descent. We used the ADAM optimization algorithm (Kingma and Ba, 2014) with a learning rate of 0.0001, L_2_ regularization factor 1.2-1.5, mini-batch size of 64, and epoch 259-519. The classifier was trained using 80% of the dataset, and the remaining 20% was held out as a validation set, with training and testing sets containing equal proportions of different stimuli. The number of iterations for the network is 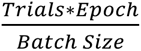, and for classification tasks, the training accuracy was validated every 20 iterations. The resulting testing accuracy was averaged over all animals and imaging sessions. For the regression task (high-resolution sAM rate decoding), the decoding error is expressed in octaves,

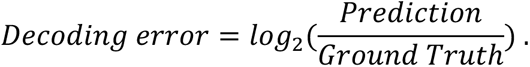

We trained the CNN decoder to perform different decoding tasks. For the first task with multiple pairs of sAM depths and -rates, we trained the decoder to classify the joint combination of sAM depths and -rates (Figure 4B), sAM rates under a given depth (Figure 4C, D), and sAM depths under a given rate (Figure 6). The classifier was further trained on an exclusive sAM depth or - rate to decode sAM rates or -depths, respectively. This trained classifier was then tested on data with other sAM depths or -rates to predict rates or -depths, and the resulting decoding accuracy was fitted by a gaussian model (Figure 7B). Validation of our model was performed by assessing the confusion matrix, feeding the classifier a shuffled neuron order in the input and receiver operating characteristic (ROC, Figure 4E). For the second experiment with more narrowly spaced sAM rates and 100% sAM depth, we trained the CNN decoder to regress, not classify, sAM rates (Figure 9D-E).

### Classifier network d-prime

To evaluate the sAM rate classification performance, we calculated the d-prime for the decoding performance of the classifier network. We define the decoder d-prime the same as the neurometric d-prime, calculating the hit rate and false alarm rate based on the confusion matrix from every imaging session. The d-prime was calculated via the one-vs-rest manner, which assumes that the positive class only includes a specific sAM rate, while the negative class contains the rest of sAM rates.

### Pattern correlation

To measure similarity of neural population activities associated with different sAM sounds, we adapted the pattern correlation described in the work of Friedrich and Laurent (2001) and population vector correlation in the work of Deitch et al. (2021) as follows. We first trial-averaged responses of single neuron associated different combinations of sAM depths and -rates. Pattern correlation of trial-averaged neural population activity in shell IC between two different sAM depths or -rates calculated on a per-frame basis using Pearson’s correlation coefficient.

### t-distribution stochastic neighbor embedding (t-SNE) clustering

The t-SNE analysis, a dimensionality reduction technique where the neural population activity is represented as a single data point in a two-dimensional map, was performed according to the framework of van der Maaten and Hinton (2008). The local proximity of data points in the map captures the similarity of data in the original dimension and t-SNE can thus cluster the data points.

In our case, the data contained N vectors: *x*_1_, …, *x_n_*, where each *x* is a neural population activity vector containing the mean ΔF/F of every neuron during sound presentation. t-SNE transforms the Euclidean distance of a pair of vectors ***x_i_*** and ***x_j_*** in the high dimensional data into a conditional probability *p*_*i*|*j*_, representing the similarity between two vectors,

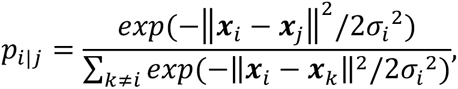

where the variance σ*_i_*^2^ is determined such that, the beforehand determined perplexity *P* satisfies

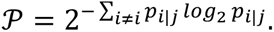

For our study, the perplexity ranges from 10 to 50. The t-SNE algorithm then constructs a mapping of a low dimensional representation ***y_i_*** and ***y_j_***, which is determined via a random walk approach, to the high dimensional data ***x_i_*** and ***x_j_***. The similarity between the low dimensional counterparts is represented by the conditional probability *q_i_*_|*j*_ of ***y_i_*** and ***y_j_*** is given by the student t-distribution with one degree of freedom,

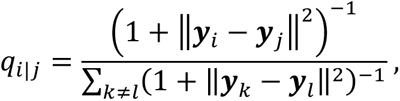

The aim of t-SNE is to minimize the mismatch between *q_i_*_|*j*_ and *p_i_*_|*j*_, so that the low dimensional counterpart of ***y_i_*** and ***y_j_*** can optimally model the similarity between the high dimensional data ***x_i_*** and ***x_j_***. Thus, the t-SNE adopts the Knullback-Leibler divergence of *q_i_*_|*j*_ and *p_i_*_|*j*_ as a loss function C,

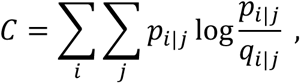

and adjusts the low dimensional representation by minimizing C via adaptive gradient descent. The optimized low dimensional counterpart ***y_1_*, …, *y_n_***, two dimensional in our case, was visualized. The values in the axes of two dimensions given by t-SNE do not have a specific meaning, but only capture the similarity in the original data.

### Statistics

Statistical assessments were performed using MATLAB_R2021b and GraphPad Prism 9. For all within- and between-group comparisons, data were tested for normality using Lilliefors’ test. Parametric approaches were adopted for data following a normal distribution, while non-parametric approaches were adopted for data not normally distributed. The alpha level is corrected for multiple comparisons using the Bonferroni-Holm method. For repeated-measures ANOVA, Geisser-Greenhouse correction was applied when sphericity is not assumed. We report mean ± SEM or median ± SEMedian 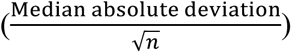for data following normal and non-normal distribution, respectively.

## Results

### Multi-photon Ca^2+^ imaging of sAM selectivity in shell IC neurons

We used a viral vector to express a genetically encoded Ca^2+^ indicator (GcaMP6f or 6s; see Methods) in the IC of mice, and then used 2-photon imaging to record neural activity in shell IC neurons as awake, head-fixed mice listened to sAM narrow-band noises (Figure 1A-C; 65-70 dB SPL, 0.5-1 s duration, 5-200 Hz sAM rate, 0-100 % sAM depth, carrier bandwidth: 16 ± 2 kHz). Imaging data were collected from fields of view (FOVs) located across the medial-lateral axis of the dorsal IC 20-55 µm from the brain surface (Figure 1C).

**Figure 1.**
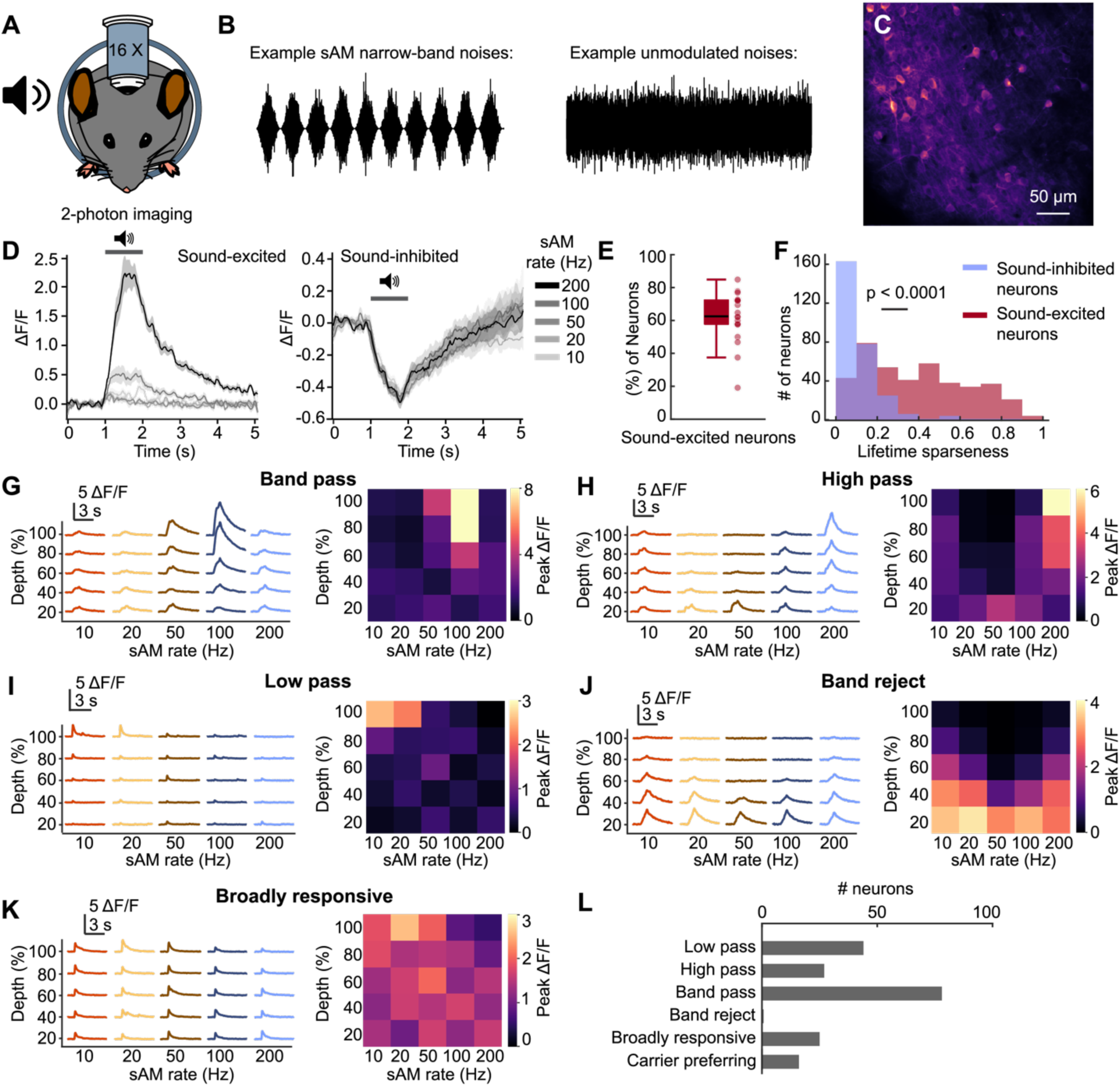
Responses of mouse shell IC neurons to sAM and unmodulated narrow-band noises. **A**. Experimental setup of sound presentation and 2-photon imaging of head-fixed, awake mice. **B**. Left, an example of presented sAM narrow-band noises (100% sAM depth and 10 Hz sAM rate). Right, an example of presented unmodulated noises. **C**. Example imaging FOV. **D**. Example responses of sound-excited (left), and sound-inhibited (right) neurons to fully modulated sAM sounds. Data are mean ± SEM. **E**. Proportion of sound-excited neurons in each imaging session. **F**. Distribution of lifetime sparseness of sound-excited and sound-inhibited neurons. A neuron is maximally selective to sAM stimuli if its sparseness is 1 and is totally unselective if the sparseness is 0. Mann-Whitney U test. **G**-**K**. Example of trial-averaged neuronal responses (left) and peak ΔF/F (right) of band pass (**G**), high pass (**H**), low pass (**I**), and band reject (**J**), broadly responsive (**K**) neurons under different sAM sounds. **L**. Distribution of the number of tuning characteristics of shell IC neurons.

We recorded a total of 1986 regions of interest (i.e., neurons) in 19 fields of view from N = 8 mice. Of these, n = 689 (34.7 %) were significantly sound-responsive as determined by a ‘signal autocorrelation’ bootstrapping analysis (Geis et al., 2011; Wong and Borst, 2019) and were used for further analysis. In agreement with classic single- and multi-unit data from central IC neurons (Rees and Møller, 1983, 1987; Krishna and Semple, 2000; Nelson and Carney, 2007), we found that n = 409 (59.4 %) sound-responsive shell IC neurons displayed a sound-evoked increase in fluorescence, i.e., they increased firing rates to sAM sounds (Figure 1D, left). Interestingly, n = 280 (40.6 %) neurons were sound-inhibited and *decreased* their fluorescence during sound presentation (Figure 1D, right); this fluorescence decrease likely reflects sound-evoked suppression of baseline firing rates (Wong and Borst, 2019).

We next quantified sAM selectivity of shell IC neurons by calculating lifetime sparseness, a measure of how discriminately a neuron responds to a set of stimuli over an observation period, for each sound-excited and sound-inhibited neuron (Rolls and Tovee, 1995; Vinje and Gallant, 2000; Willmore and Tolhurst, 2001; Kato et al., 2017). Lifetime sparseness values range from 0 to 1, reflecting the extent to which a neuron’s sound-evoked fluorescence change is unique to a single sAM sound (sparseness = 1) or alternatively, all sounds tested (sparseness = 0). Across the population, sound-excited neurons showed a larger spread of lifetime sparseness values, and thus greater selectivity to specific sAM rates compared to sound-inhibited neurons (Figure 1F; U = 182294, z = 16.051, p < 0.0001, Mann-Whitney U test). Indeed, sound-inhibited neurons were often broadly inhibited by all sounds tested, whereas sound-excited neurons typically responded maximally to a specific tested sAM rate (see examples in Figure 1D). Consequently, many sound-excited shell IC neurons displayed the canonical modulation transfer functions previously observed in the central IC (Langner and Schreiner, 1988; Krishna and Semple, 2000; Nelson and Carney, 2007): Low-pass, high-pass, band-pass, and band-reject (Figure 1G-L). The peak fluorescence response to preferred sAM rate generally increased as a function of sAM depth (Figure 1G-K, right panels), indicating that higher depths drove stronger firing rates. Altogether these data show that modulation transfer functions previously observed in the central IC are also found in the shell IC, where the neuronal response to sAM sounds increases as its depth increases. However, many neurons often had significant Ca^2+^ fluorescence responses even at non-preferred sAM rates, in agreement with the previously reported broad and variable feature selectivity of non-lemniscal IC neurons (Aitkin et al., 1975; Ito et al., 2014; Parras et al., 2017; Wang and Borst, 2019; Chen and Song, 2019).

### GABAergic and glutamatergic neurons in shell IC exhibit similar responses to sAM sounds

The IC contains two neurochemically distinct neuron classes that release either the excitatory or inhibitory neurotransmitters glutamate or GABA, respectively. However, the extent to which these distinct neurons show unique sound responses is less clear (Oliver et al., 1994; Ito et al., 2011; Ito and Oliver, 2012; Wong and Borst, 2019; Ito, 2020). To test whether sAM tuning differed between GABAergic and glutamatergic neurons, we expressed the red fluorescent protein tdTomato in GABAergic neurons in a subset of mice by crossing VGAT-ires-cre and Ai14 reporter mice (N = 4) and compared GcaMP6f responses in tdTomato-positive and - negative neurons (GABAergic and presumptive glutamatergic neurons, respectively; Figure 2A, B). GABAergic neurons comprise 11.5 % (n = 57) of the neurons in these datasets (Figure 2C). In agreement with prior studies suggesting that GABAergic and glutamatergic IC neurons exhibit largely overlapping sound responses (Ono et al., 2017; Wong and Borst 2019; Chen and Song 2019), we found that, within GABAergic neurons, 43.8 % were sound-excited and 56.2 % were sound-inhibited (Figure 2D, E), the sound-excited GABAergic and glutamatergic neurons showed qualitatively similar distributions of sAM selectivity and modulation transfer functions (Figure 2F). These results further support a limited correlation between neurotransmitter phenotype and acoustic responses in IC neurons. Given the lack of overt differences between GABAergic and presumptive glutamatergic IC neurons in their responses to sAM sounds, data from both cell-types were pooled for all subsequent analyses.

**Figure 2.**
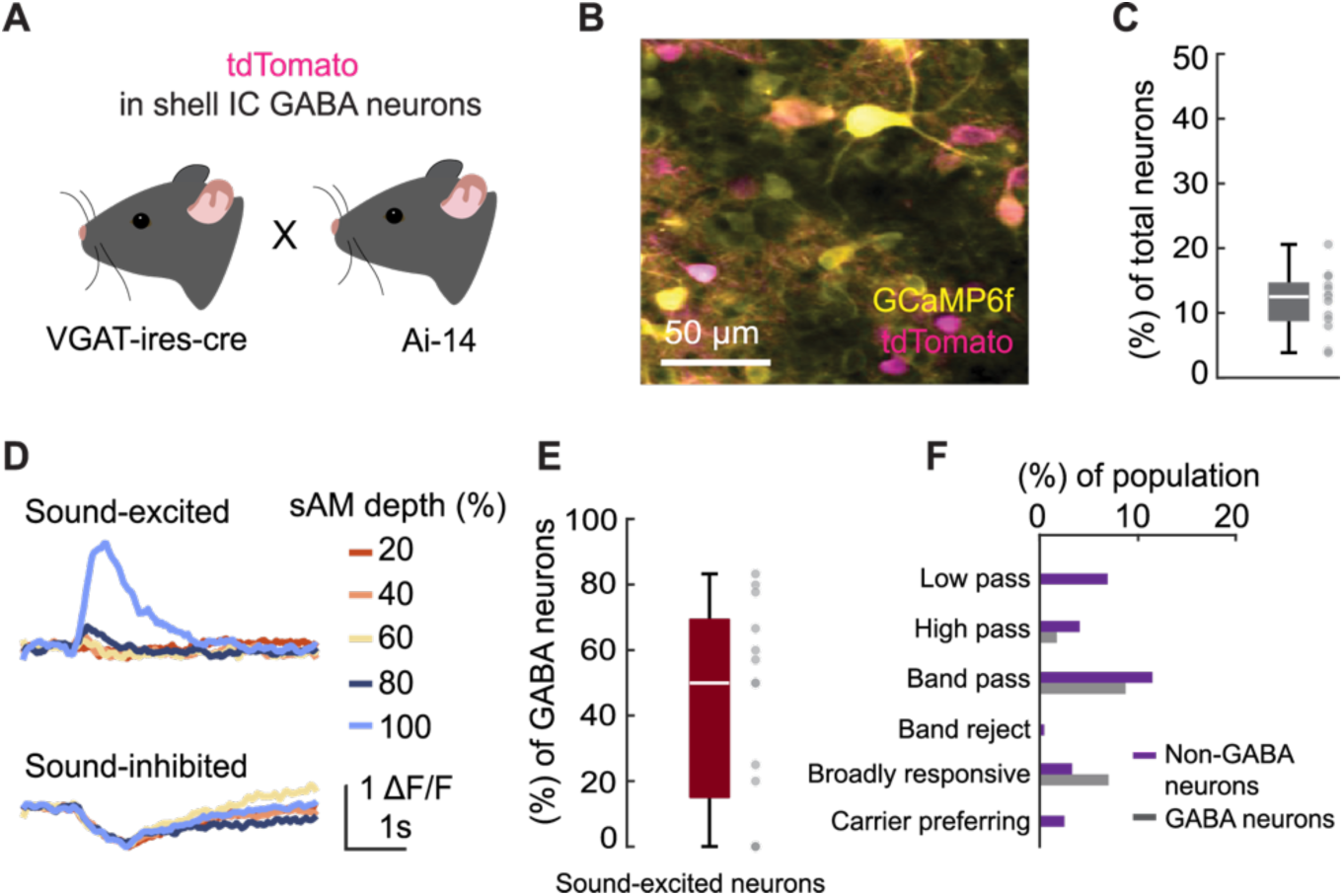
sAM tuning of shell IC GABAergic and non-GABAergic neurons. **A**. tdTomato was expressed in GABAergic neurons in transgenic VGAT-ires-cre x Ai14 mice. **B**. Example imaging FOV. **C**. Proportion of GABAergic neurons in each imaging session. **D**. Examples of sound-excited and sound-inhibited responses of GABAergic neurons to sounds with 100 Hz sAM rate and different sAM depths. **E**. Proportion of sound-excited GABAergic neurons in each imaging session. **F**. Distribution of tuning characteristics of non-GABAergic and GABAergic neurons.

### Individual shell IC neurons transmit unreliable sAM information

We further quantified sAM sensitivity by testing how shell IC neurons respond at their preferred-compared to their non-preferred sAM rate. To this end, we calculated the neurometric sensitivity index (d-prime, von Trapp et al., 2016) for 668 sAM preferring neurons, counting responses at the preferred sAM rate as a hit, whereas responses at other sAM rates were counted as false alarms (Figure 3A). Sound-excited neurons had a higher neurometric d-prime compared to sound-inhibited neurons (Figure 3B; d-prime for sound-excited and inhibited neurons at 100% sAM depth: 0.96 ± 0.03 and 0.20 ± 0.02, respectively. Main effect of sAM depth: F (4,16690) = 197.6, p < 0.0001. Main effect of neuron type: F (1,16690) = 912.4, p < 0.0001. Interaction: F (4, 16690) = 124.1, p < 0.0001, two-way ANOVA). These results qualitatively mirror our previously observed differences in lifetime sparseness (e.g., Figure 1F). When averaged across the entire neuronal population, the mean d-prime increased monotonically as a function of the sAM depth for the preferred sAM rate, while changing the sAM depth of non-preferred sAM rates had little impact on the d-prime (Figure 3C, left). However, this effect was primarily driven by n = 89 (13 %) sound-excited neurons whose d-prime values increased sharply as a function of sAM depth (Figure 3B), and only a small population n = 15 (2 %) of sound-excited neurons showed the opposite trend, i.e., neurometric discrimination being inversely proportional to sAM depth (Figure 3C). This result may reflect the fact that in sound-excited neurons, increasing sAM depth preferentially increased fluorescence responses at the neuron’s preferred sAM rate, whereas increasing sAM depth of non-preferred sAM rates had minimal effect on neuronal activity (Figure 3D; Main effect of sAM depth: F (1.488, 598.1) = 44.48, p < 0.0001. Main effect of preference: F (1, 402) = 99.79, p < 0.0001. Interaction: F (1.708, 686.8) = 60.73, p < 0.0001, two-way repeated-measures ANOVA). Altogether, these data show that a subset of shell IC neurons can reliably discriminate sAM rates, in a sAM depth-dependent manner. However, most individual neurons still displayed poor d-prime profiles especially at low sAM depths (Figure 3B), suggesting that many shell IC neurons transmit variable, and potentially unreliable, information regarding sAM features.

**Figure 3.**
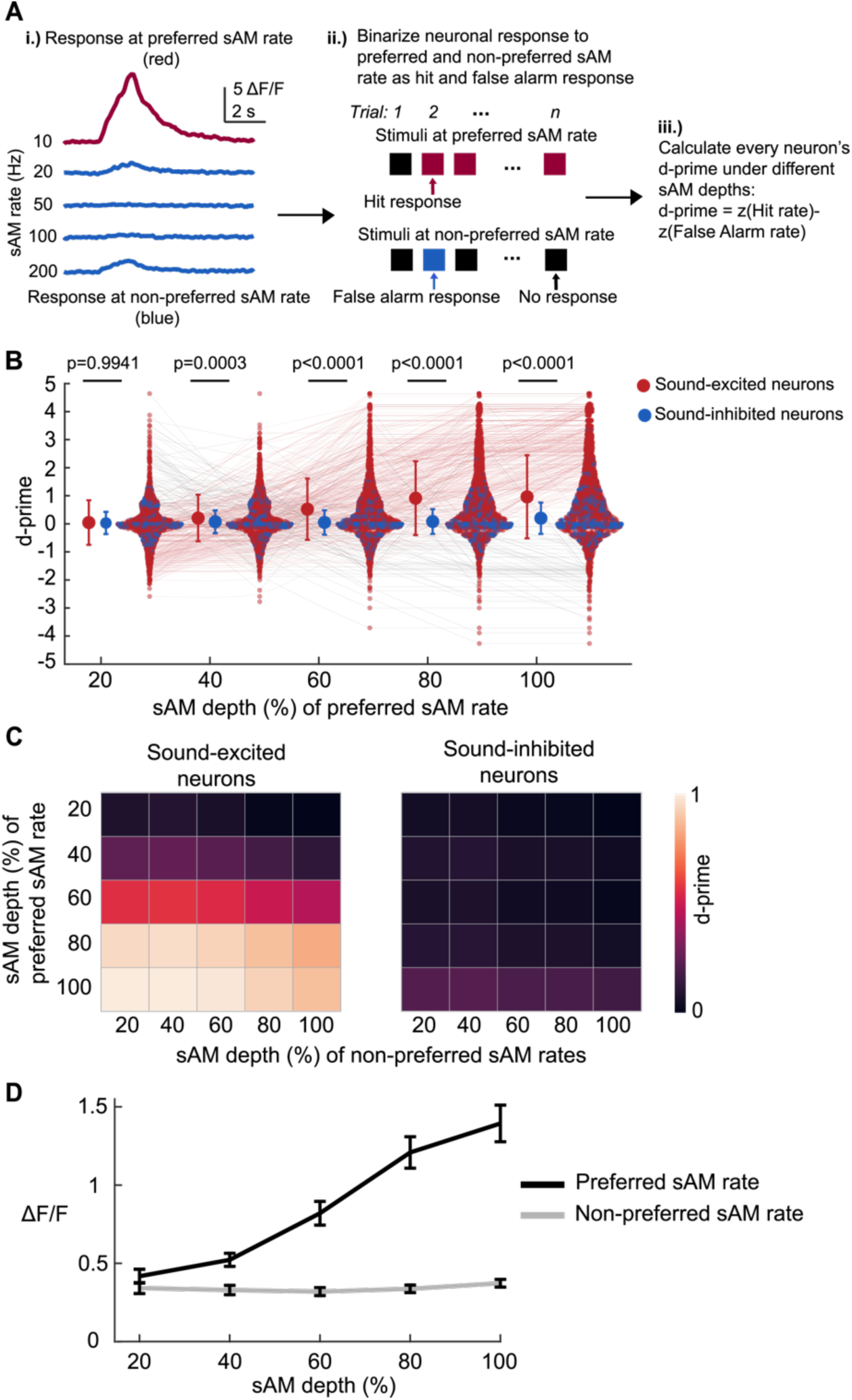
Neurometric sensitivity of individual shell IC neurons. **A**. Schematic of neurometric sensitivity index (d-prime) analysis. Preferred sAM rate is determined by the peak fluorescence response at different sAM rates (i). We binarize a neuron’s response as a hit if the mean neuronal response at its preferred sAM rate exceeds three times the standard deviation of the baseline fluctuation on a single trial. Similarly, we count a false alarm response if a neuron’s average response at its non-preferred sAM rate exceeds three times the standard deviation of the baseline fluctuation (ii). The d-prime was then calculated for each neuron for all pairs of sAM depths for preferred and non-preferred sAM rates (iii). **B**. Distribution of d-prime at varying sAM depths for the preferred sAM rate. Each line indicates a neuron displaying an increasing (pink) or decreasing (gray) trend across sAM depths for the preferred sAM rate (See Methods). Data are mean with ± std. Two-way ANOVA. Šídák’s multiple comparison between d-primes of sound-excited and sound-inhibited neurons. **C**. Left, averaged d-prime of sound-excited neurons. Right, averaged d-prime of sound-inhibited neurons. **D**. Trial-averaged fluorescence response to preferred and non-preferred sAM rate. Data are mean with ± SEM.

### Neural population coding of sAM rate

Our data suggest that single neuron discrimination of sAM rate is variable, particularly at low sAM depths. However, in circuits where the feature selectivity of individual neurons is noisy or ambiguous, neural population codes (i.e., collective activity in large sets of neurons) nevertheless provide an accurate representation of sensory signals for downstream circuits (Partridge et al., 1981; Lee et al., 1988; Vogels, 1990; Heiligenberg, 1991; Stringer et al, 2021; Robotka et al., 2023).

We thus tested the extent to which sAM features are encoded in the activity of shell IC neuronal *populations*. While the pattern in population activity is hard to interpret mechanistically, machine learning algorithms are valuable tools to extract meaningful information from neural population codes. To identify neural population response pattern to sAM sounds while preserving the temporal fidelity of the neural data, we employed a contemporary artificial convolutional neural network (CNN), similar to previous approaches applied to visual and auditory systems (Anumanchipalli et al., 2019; Sun et al., 2020; Zhang et al., 2020).

We trained the CNN to decode sAM features using time series fluorescence responses from all significantly sound-responsive shell IC neurons in our datasets (Figure 4A). We first tested if the CNN could jointly classify the sAM depth and -rate of the sound on a trial-by-trial basis. Accordingly, joint decoding accuracy significantly exceeded chance level (W = 190, p = 0.0001, Wilcoxon signed-rank test), and the sAM rate was classified more accurately at higher depths (Figure 4B). These data suggest that the representational fidelity of sAM rate in shell IC populations increases with sAM depth, such that a CNN’s sAM rate classification accuracy should vary with the sAM depth of its training data. To test this hypothesis, we next trained CNNs on data from all sAM rates under a specific sAM depth, and quantified rate decoding accuracy at each depth. Accuracy was not significantly different from chance level when CNNs were trained on 20 % sAM depth (Figure 4C). However, sAM rate classification accuracy increased sharply above chance level when CNNs were trained with higher sAM depths, peaking at 81 % ± 4 % when trained on fully modulated sAM sounds (Figure 4C, D. Main effect of sAM depth: F (2.496, 44.92) = 17.83, p < 0.0001. Main effect of treatment group (trained vs. chance level): F (1,18) = 127.7, p < 0.0001. Interaction: F (2.517, 45.31) = 25.92, p < 0.0001, two-way repeated-measures ANOVA). Moreover, classification errors tended to cluster along the diagonal of confusion matrices (Figure 4C), implying that the population representation of sAM rates varies along a continuum. Importantly, classification accuracy was relatively impervious to changes in hyperparameters such as the kernel size in the convolutional layer (*X*^2^ (4) = 6.42, p = 0.44, Friedman test for sAM rate decoding accuracy under 100% depth with different kernel size). Moreover, shuffling neurons in population response matrices had little impact on the decoding performance of sAM rate classification (W = 42.5, p = 0.5571, Wilcoxon signed-rank test for sAM rate decoding accuracy under 100% depth when neuron is shuffled and control). Altogether, these results suggest that sAM rate representation in the shell IC is supported by a population code such that representational fidelity is steeply dependent on sAM depth. These results are in qualitative agreement with perceptual thresholds determined from behaving rodents engaged in AM detection tasks (Kelly et al., 2006; Caras and Sanes, 2017; van den Berg et al., 2023).

**Figure 4.**
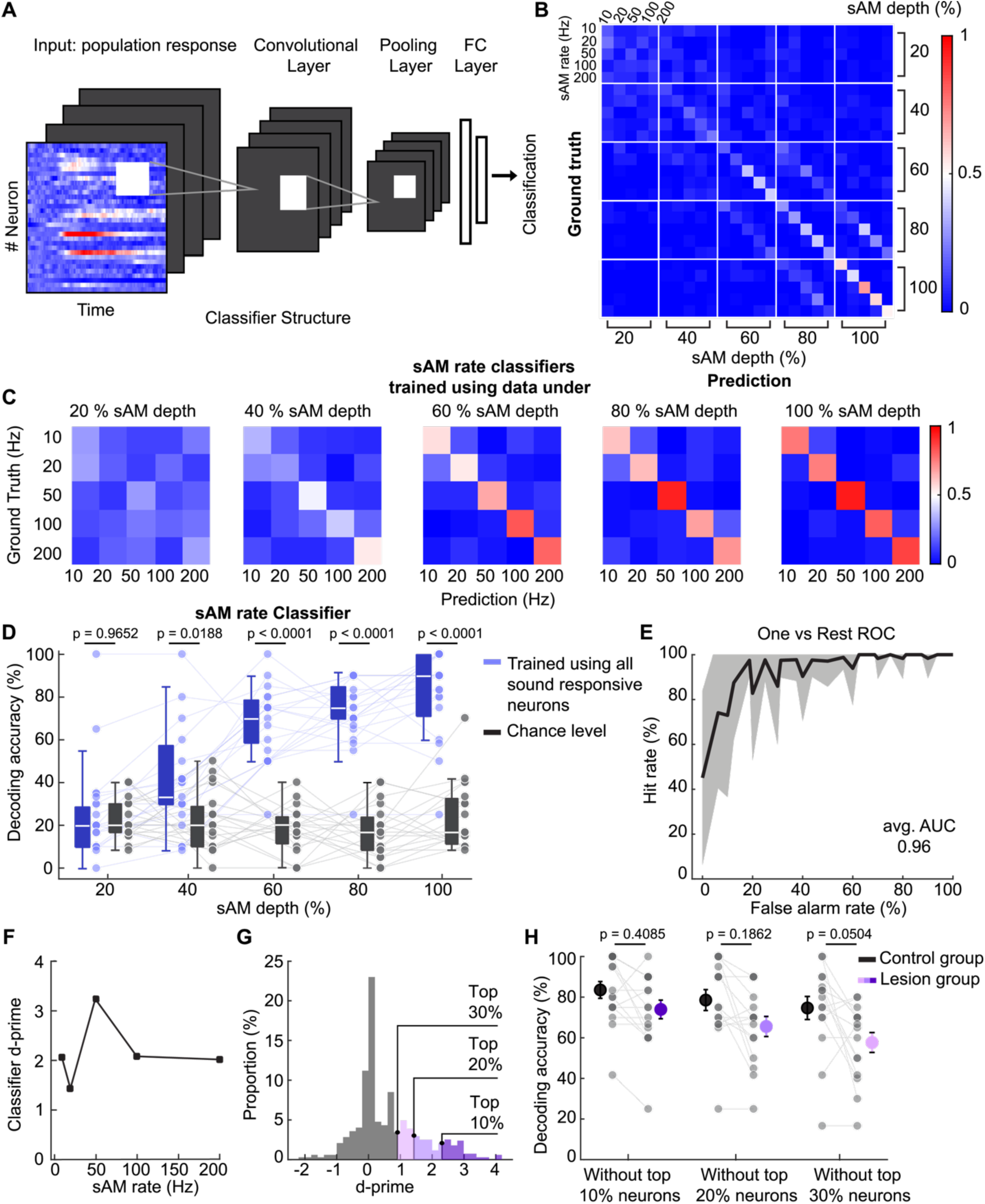
Decoding sAM features using shell IC neural population activity. **A**. Structure of the CNN classifier. A CNN classifier fed with Ca^2+^ signal time series was trained to classify sAM features. **B**. Normalized confusion matrix of sAM depth and -rate joint combination classification averaged across imaging sessions. **C**. Normalized confusion matrix of sAM rate classification under a given sAM depth averaged across imaging sessions. **D**. Decoding accuracy of the sAM rate classifier under a given depth and the corresponding chance level. Two-way repeated-measures ANOVA. Šídák’s multiple comparison between the chance level accuracy and decoding accuracy trained using all neurons. **E**. One-vs-rest ROC of sAM rate classification under 100% sAM depth. Data are mean with ± std. **F**. d-prime of sAM rate classifier under 100% sAM depth. **G**. Individual neuron d-prime distribution (under 100 % sAM depth), with the corresponding highly sensitive neurons falling within top deciles (10 %, 20 %, 30 % of d-prime). **H**. Decoding accuracy of the sAM rate classifier under 100% sAM depth, trained while excluding the top deciles (10%, 20%, 30%) of neurons, selected based on their d-prime profiles in the left panel. To rectify the effect of the number of neurons on the decoding performance of the classifier, decoding accuracy of the classifier trained with a balanced number of randomly chosen neurons was visualized as control. Data are mean with ± SEM and gray dots represent predictions from each imaging session. Two-way ANOVA. Šídák’s multiple comparison between the decoding accuracy trained without highly tuned neurons and the control.

### Noisy population codes provide reliable estimates of sAM features

The CNN model’s high performance may reflect a robust capacity for sAM rate discrimination despite an otherwise low sensitivity at level of individual shell IC neurons. In agreement, the average area under receiver operating characteristic (ROC) curve (AUC) of the sAM rate classifier under 100 % sAM depth reached 0.96 (Figure 4E). We further tested the classifier’s discrimination ability by calculating a d-prime for sAM rate classification under 100 % depth. The CNN classifier achieved a mean d-prime of 2.16 ± 0.29, more than 3-fold higher than the mean neurometric d-prime for single neurons at 100 % sAM depth (0.66 ± 0.02; Figure 4F). Importantly, this result did not simply reflect the classifier’s reliance on a minority of shell IC neurons that are highly tuned to specific sAM rates: We conducted an *in silico* “lesion” experiment where we trained the sAM rate decoder while excluding from training datasets highly tuned neurons, as characterized by top deciles of d-prime (10 %, 20 %, and 30 %; Figure 4G). We then tested the extent to which sAM rate classification accuracy deteriorates relative to control training conditions. We minimized the effect of neuron count on classifier accuracy by randomly excluding the same number of neurons from control classifiers as removed from the “lesion” condition (Averbeck et al., 2006; Yoshida and Ohki, 2020; Zhang et al, 2020; Stringer et al., 2021). Unsurprisingly, the mean decoding accuracy of the sAM rate classifier was reduced when trained without highly tuned neurons, although this reduction was modest: “Lesioned” classifiers nevertheless achieved a greater than three-fold greater accuracy above chance level (Figure 4H, 74 % ± 5 %, 66 % ± 5 %, and 58 % ± 5 % for removal of 10, 20, and 30 % top deciles respectively. A two-way ANOVA revealed a main effect of treatment group (i.e. lesion vs. control: F (1, 86) = 10.91, p = 0.0014) and a main effect of lesion size (F (2, 86) = 3.353, p = 0.0398) but no significant interaction (F (2, 86) = 0.2909, p = 0.7483), and Šídák’s multiple comparison tests revealed no statistical significance between lesion and control groups across all lesion sizes (Figure 4H). Thus, we conclude that sAM rate identity is robustly represented via a population code, despite a noisy and otherwise low neurometric discrimination measured in individual neurons.

### Sound-inhibited neurons contribute minimally to sAM population codes

Our training datasets thus far included all sound-responsive neurons regardless of whether they were excited or inhibited by sAM sounds. However, our results of Figure 1F show that sound-inhibited neurons generally exhibit low lifetime sparseness and thereby are broadly tuned to sAM rates. To what extent do sound-inhibited neurons contribute to sAM rate representations? We first tested this idea by removing sound-excited neurons from the training datasets, such that sAM rate classifier training progressed exclusively using sound-inhibited neurons. Interestingly, classifiers trained under these conditions showed decoding accuracies that differed significantly from chance-level. However, the summary data lacked the strong relationship between sAM depth and decoding accuracy observed in Figure 4D, where classifiers were trained on the entire neuronal population (Figure 5A; Main effect of sAM depth: F (2.9, 46.4) = 1.023, p = 0.3891. Main effect of treatment group: F (1, 16) = 14.36, p = 0.0016. Interaction: F (3.382, 54.12) = 4.725, p = 0.0039, two-way repeated-measures ANOVA). Consequently, classifiers trained exclusively on sound-inhibited neurons exceeded chance level only when trained with 60% and 100% depth data (Figure 5A, Šídák’s multiple comparison tests), which qualitatively aligns with our analyses showing mostly uniform sound-evoked inhibition in shell IC neurons (Figure 1).

**Figure 5.**
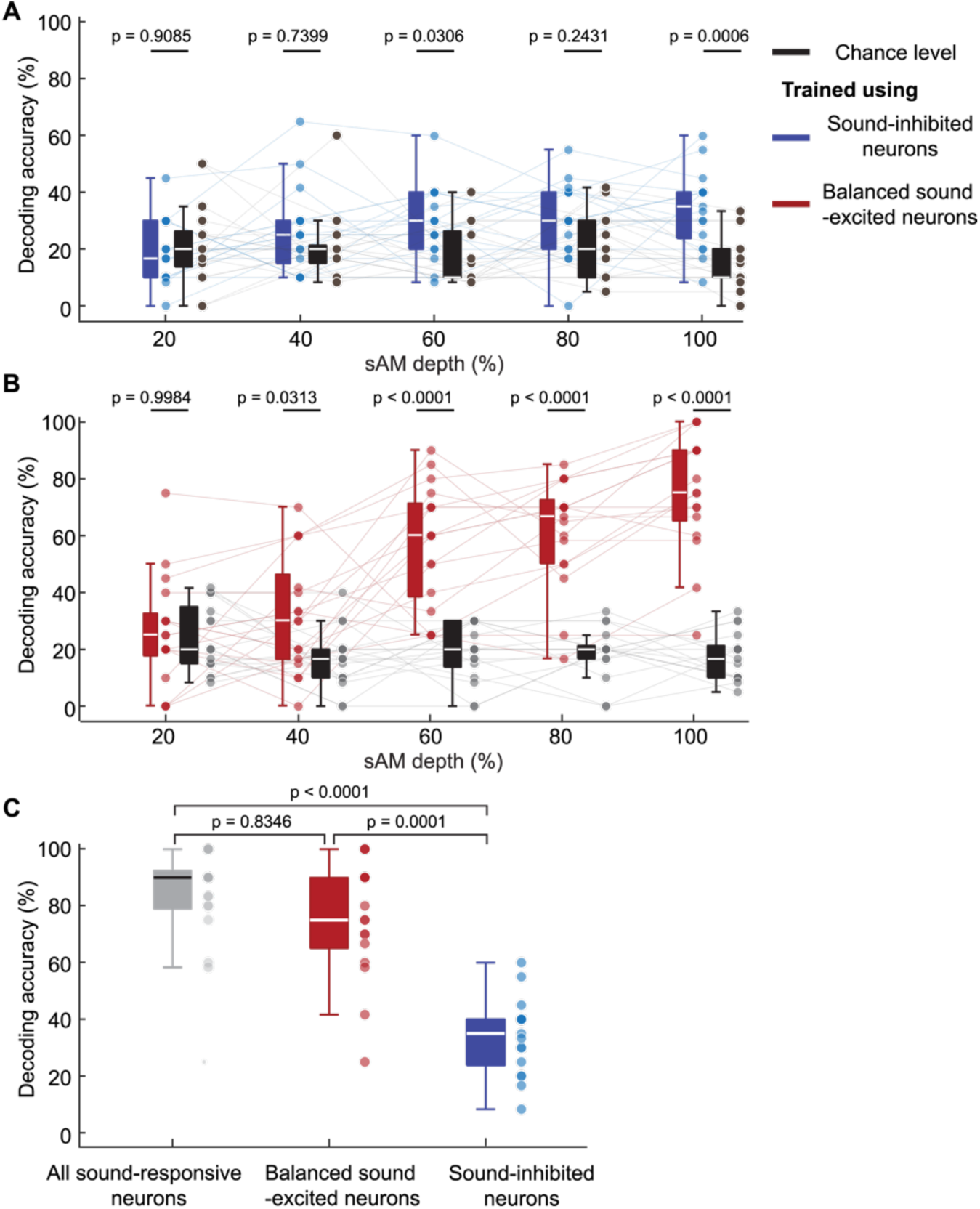
Decoding sAM rate using sound-excited and sound-inhibited neurons. **A**. Decoding accuracy of sAM rate classifier trained using sound-inhibited neurons and corresponding chance level. Two-way repeated-measures ANOVA. Šídák’s multiple comparison between decoding accuracy trained using sound-inhibited and chance **B**. Decoding accuracy of sAM rate classifier trained using sound-excited neurons and corresponding chance level. To ensure an equal number of sound-excited and sound-inhibited neurons, we randomly selected a subset of sound-excited neurons from each imaging session. Two-way repeated-measures ANOVA. Šídák’s multiple comparison between decoding accuracy trained using balanced sound-excited and chance. **C**. Decoding accuracy of sAM rate classifier trained using all sound-responsive, sound-excited, and sound-inhibited neurons under 100 % sAM depth. Kruskal-Wallis test. Dunn’s multiple comparison between decoding accuracies trained under different conditions.

We next tested how removing sound-inhibited neurons from the training datasets impacts sAM rate decoding accuracy. To ensure that any result does not reflect a spurious effect of imbalanced sound excited and inhibited neuron counts (n = 409 and n = 280, respectively; Figure 1), we randomly removed subgroups of sound-excited neurons such that the total neuron number in each training session was the same as for classifiers trained exclusively with sound-inhibited neurons. CNNs trained on balanced sound-excited neurons recapitulated the depth-dependent increase in classification accuracy observed in Figure 4D (Figure 5B; Main effect of sAM depth: F (2.637, 42.2) = 16.06, p < 0.0001. Main effect of treatment group (trained vs. chance level): F (1, 16) = 101, p < 0.0001. Main effect of interaction: F (2.573, 41.16) = 20.25, p < 0.0001, two-way repeated-measures ANOVA). Consequently, sAM rate classifiers under 100 % depth trained using balanced sound-excited neurons performed significantly higher than classifiers trained using only sound-inhibited neurons, and had accuracies comparably to classifiers trained using all sound-responsive neurons (Figure 5C; H (2) = 27.4074, p < 0.0001, Kruskal-Wallis test; See Dunn’s multiple comparison tests). These results indicate that activity of sound-excited neurons suffices to represent sAM sound identity in the shell IC, and align with our observation that sound-inhibited neurons are generally suppressed by all sAM stimuli (Figure 1F). Thus, sound-evoked firing rate increases likely transmit the bulk of informative shell IC efferent signals. More broadly, these conclusions agree with recent data from the primary visual cortex suggesting that increases, rather than decreases in neuronal firing rate are the dominant source of perceptually relevant neural activity (Cone et al., 2020).

### CNNs classify absolute sAM depth

Detection thresholds for sAM sounds can depend on sAM rate in rodents, birds and humans (Dooling and Searcy, 1981; Bacon and Viemeister, 1985; Klump and Okanoya, 1991; van der Berg et al., 2023). We therefore asked if population representations of sAM depth are differentially informative depending on sAM rate. To this end, we iteratively trained CNN classifiers on data from all sAM depths under a specific sAM rate and tested if sAM depth classification accuracy varied as a function of the training sets’ sAM rate. Similar to our analysis of joint rate- and depth decoding accuracy, CNNs classified the absolute sAM depth significantly above chance for all training sets, although accuracy was only marginally (and not significantly) dependent on sAM rate (Figure 6A, B. Main effect of sAM rate: F (3.4, 61.21) = 2.396, p = 0.0694. Main effect of treatment group (trained vs. chance level): F (1, 18) = 202.8, p < 0.0001. Interaction: F (2.942, 52.96) = 3.789, p = 0.016, two-way repeated-measures ANOVA). Moreover, confusion matrices showed that classification errors were most likely to occur between neighboring sAM depths, similar to the results observed in psychometric sAM depth discrimination (Figure 6B; Lee and Bacon, 1997). Our results thus show that sAM depth is reasonably well represented in the shell IC, and that this discriminative ability remains robust over an order of magnitude difference in sAM rate.

**Figure 6.**
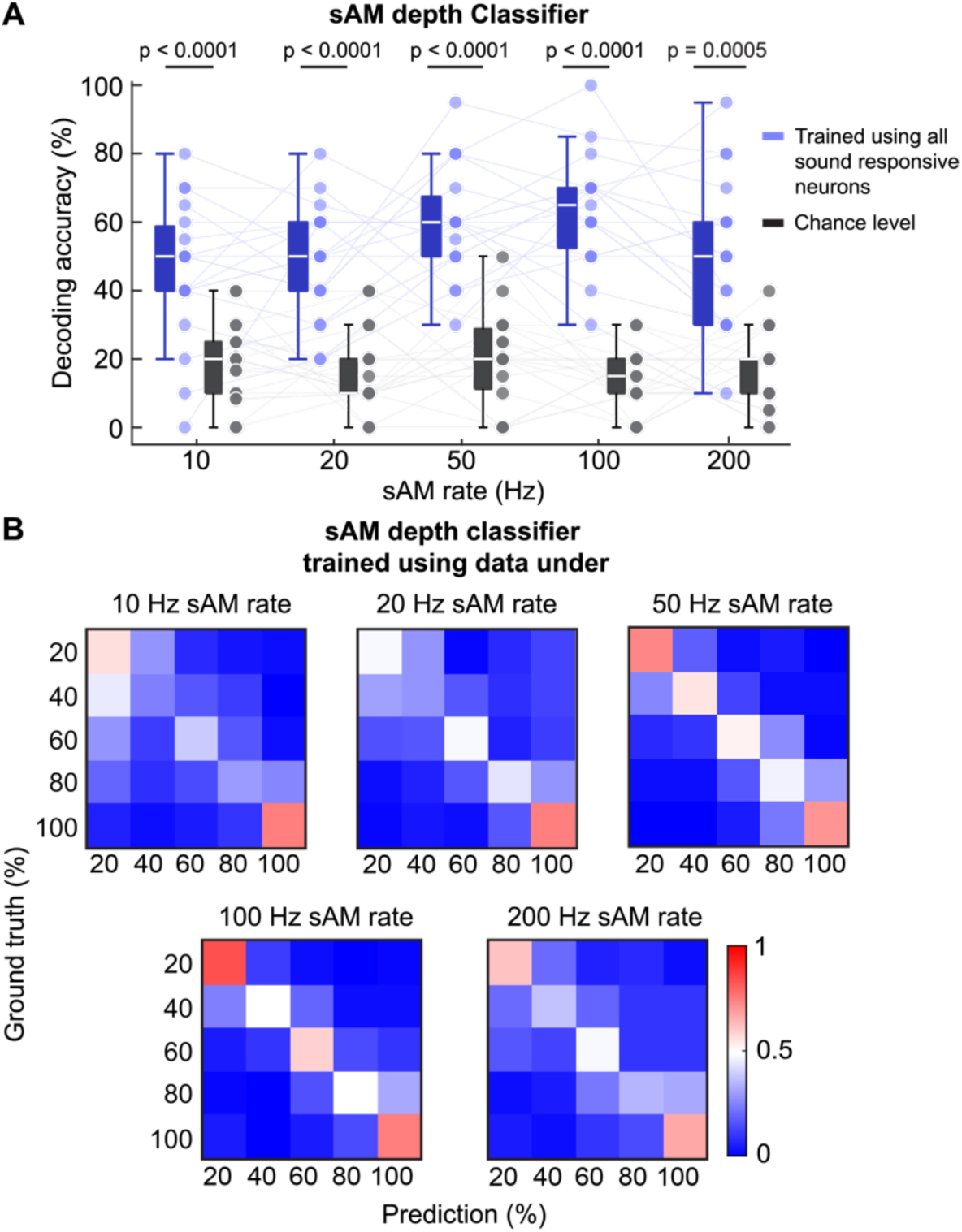
Decoding sAM depth. **A**. Decoding accuracy of the sAM depth classifier under a given rate and its corresponding chance level. Two-way repeated-measures ANOVA. Šídák’s multiple comparison between decoding accuracy trained using all neurons and the chance. **B**. Normalized confusion matrix of sAM depth classification under a given rate averaged across imaging sessions.

### sAM representation diverges across different sAM depths

sAM rate classification accuracy depended steeply on the sAM depth of the training set and saturated near 60-80 % depth (Figure 4D). These results suggest that representational fidelity increases monotonically with the depth of the sAM sounds. If this hypothesis is correct, a CNN classifier trained under a single sAM depth might still decode the rate of stimuli with lower or higher sAM depths at above chance accuracy. However, this hypothesis further predicts that decoding accuracy under these conditions should vary as a function of the depth difference between training and testing datasets.

To this end, we trained CNN classifiers on data from all sAM rates under a single sAM depth. We then measured classification accuracy on both original testing sets held back from training (e.g., datasets with the same sAM depth as the training set data), as well as extended testing sets consisting of data from lower or higher sAM depths not incorporated in training (Figure 7A). With the exception of CNNs trained on 20 % sAM depth, classification accuracy remained above chance when evaluating held-out data of the same depth (Figure 7B, see also Figure 4D. 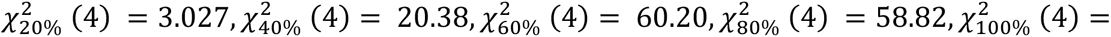 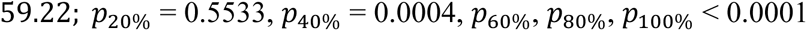, Friedman test for each sub-panel in B). Interestingly, CNN performance was capped by the training set’s sAM depth: When tested on extended datasets comprised of higher sAM depths than the training set, decoding accuracy rarely exceeded the accuracy of the held-out data (Figure 7B; see results of post-hoc tests). Furthermore, decoding accuracy decreased monotonically instead of randomly as a function of distance between training and testing sAM depths when the extended datasets’ sAM depth was lower than that of training data; this effect is particularly striking when the model is trained at 80 % and 100 % depths (Figure 7B). One potential interpretation is that population responses to different sAM rates become increasingly dissimilar at higher sAM depths. Additionally, increasing sAM depth likely drives higher firing rates in shell IC neurons, thereby increasing the amplitude of Ca^2+^ signals which reflect mean firing rates in a specific time window. Either of these scenarios could account for the observation that classifiers trained on high depths can robustly classify sAM rate at lower depths, whereas increasing testing sAM depth has little effect when the model was exclusively trained on low sAM depths (Figure 7B).

**Figure 7.**
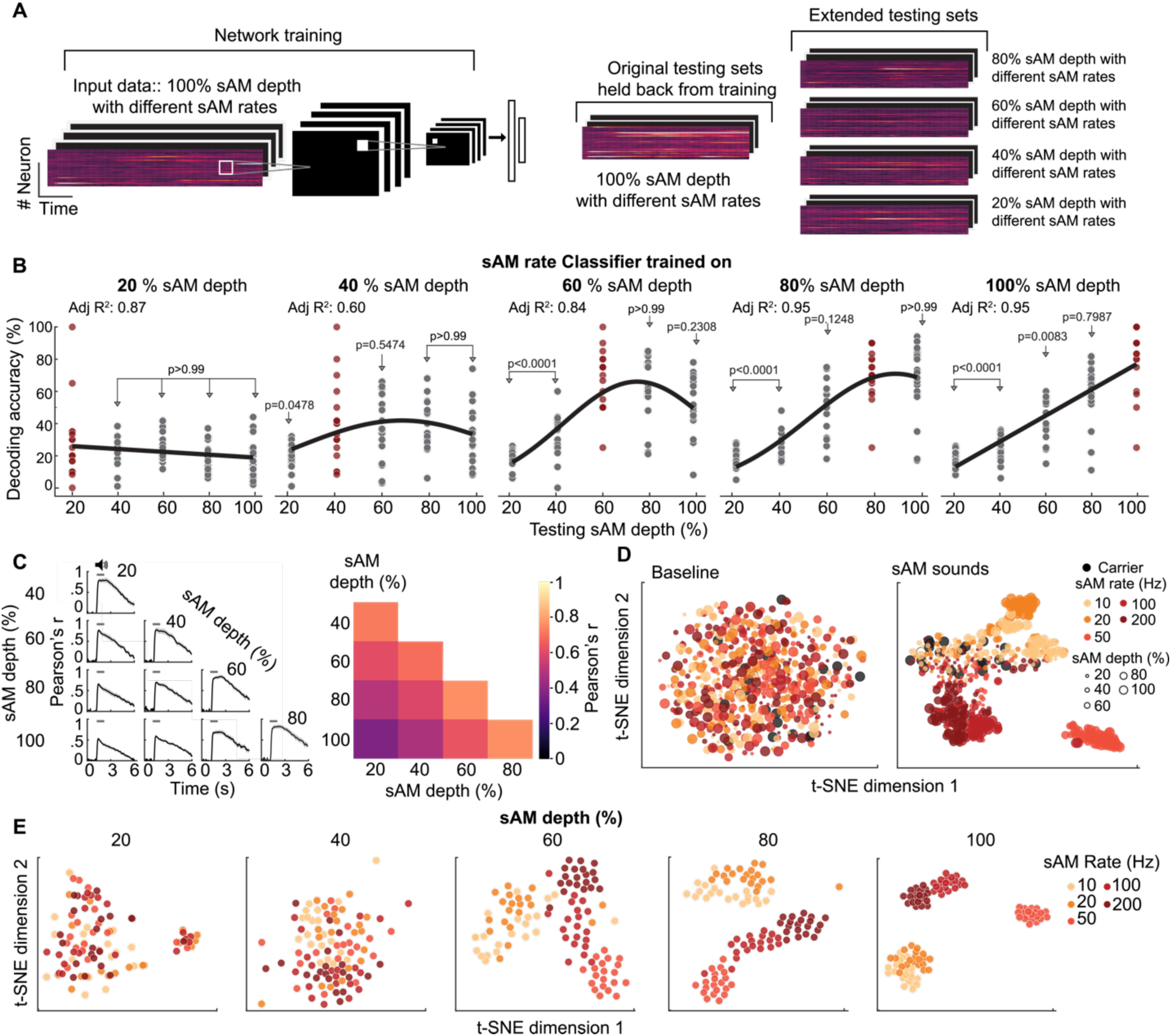
Representation of different sAM depths in the shell IC. **A**. For sAM rate classification, the decoder was trained using input data from a specific sAM depth. After training was complete, the decoder was evaluated on both original testing sets held back from training and extended testing sets (all datasets from other sAM depths). **B**. Evaluation of the sAM rate classifier on extended testing sets (black) and held-back testing sets (red). The curve was fitted using a gaussian model. Friedman test for each sub-panel. Dunn’s multiple comparison between decoding accuracies tested on held-back and extended testing sets. **C**. Pattern correlation of each pair of sAM depths. Left, Correlation of the trial-averaged neural population vector across two different sAM depths or -rates on a per-frame basis. Data are mean ± SEM Right, averaged correlation data during sound presentation. **D**. Two-dimensional t-SNE map of shell IC neural population activity. Left, t-SNE map of neural population activity during baseline. Right, t-SNE map of neural population activity during sound presentation. **E**. t-SNE map of neural population data with a single sAM depth only. Each dot represents a single trial neural population activity.

We further quantified the extent of similarity between different sAM depths using pattern correlation (Friedrich and Laurent, 2001; Deitch et al., 2021; see Methods). The averaged correlation data during sound presentation reveal an increased correlation as a function of sAM depth similarity (Figure 7C, right, heatmaps). These data further argue that under passive listening conditions, sAM depth appears to be represented in the shell IC population activity as a continuum, rather than as distinct categories. This similarity is also reflected in the low-dimensional representation of neural population data by applying a t-distributed stochastic neighbor embedding (t-SNE) clustering analysis (van der Maaten and Hinton, 2008), which reduces dimensionality while preserving the similarity of data in the original dimensions. The t-SNE map constructed from baseline data 1 s prior to sound onset is unstructured (Figure 7D, left), whereas the map generated from data points during sound presentation results in distinct clusters (Figure 7D, right). Instead of forming isolated islands, the sound data are spatially clustered and interconnected based on their similarity in sAM depth. Interestingly, the t-SNE data points were also separated based on sAM rate (Figure 7D, note the colors). However, this separation along the rate dimension was dictated by sAM depth: When performing separate t-SNE analyses using data from single sAM depths, clusters representing specific sAM rates increasingly diverged in a depth-dependent manner (Figure 7E).

As a separate test of how sAM rate population codes vary with sAM depth, we determined CNNs’ sAM depth thresholds by training the models to perform a binary classification task reporting the absence or presence of sAM in a band-pass noise carrier (Figure 8A). Similar to our results with CNNs classifying the absolute sAM depth (Figure 5), sAM detection accuracy increased as a function of the sAM depth and was marginally dependent on sAM rate (Figure 8B-C; Main effect of sAM depth: F (2.196, 37.33) = 30.3, p < 0.0001. Main effect of sAM rate: F (2.814, 47.84) = 4.54, p = 0.008. Interaction: F (5.755, 97.83) = 2.03, p = 0.0715, two-way repeated-measures ANOVA). The somewhat lower classification accuracy at slower sAM rates is not surprising, as this results qualitatively mirrors the observation that mice’s sAM detection thresholds are slightly increased at slower sAM rates (van den Berg et al., 2023). Additionally, a logistic function fit the mean detection accuracy across all sAM rates revealed that sAM detection threshold in this binary classification task is ∼ 52 % sAM depth (Figure 8C). Finally, we also observed that the pattern correlation between unmodulated carrier and sAM sounds decreased monotonically as a function of sAM depth (Figure 8D), similar to our results correlating fluorescence data of different sAM depth trials (Figure 7C). Altogether, the data suggest that increasing sAM depth dictates the intensity of neuronal firing rates, whereas sAM rate is represented by a combinatorial code of distinct neurons firing to subsets of sAM sounds. Consequently, tuning to sAM rate at the population level remains largely unchanged across different sAM depths at suprathreshold levels. On the other hand, increasing sAM depth sharpens the distinction between representations of different sAM rates.

**Figure 8.**
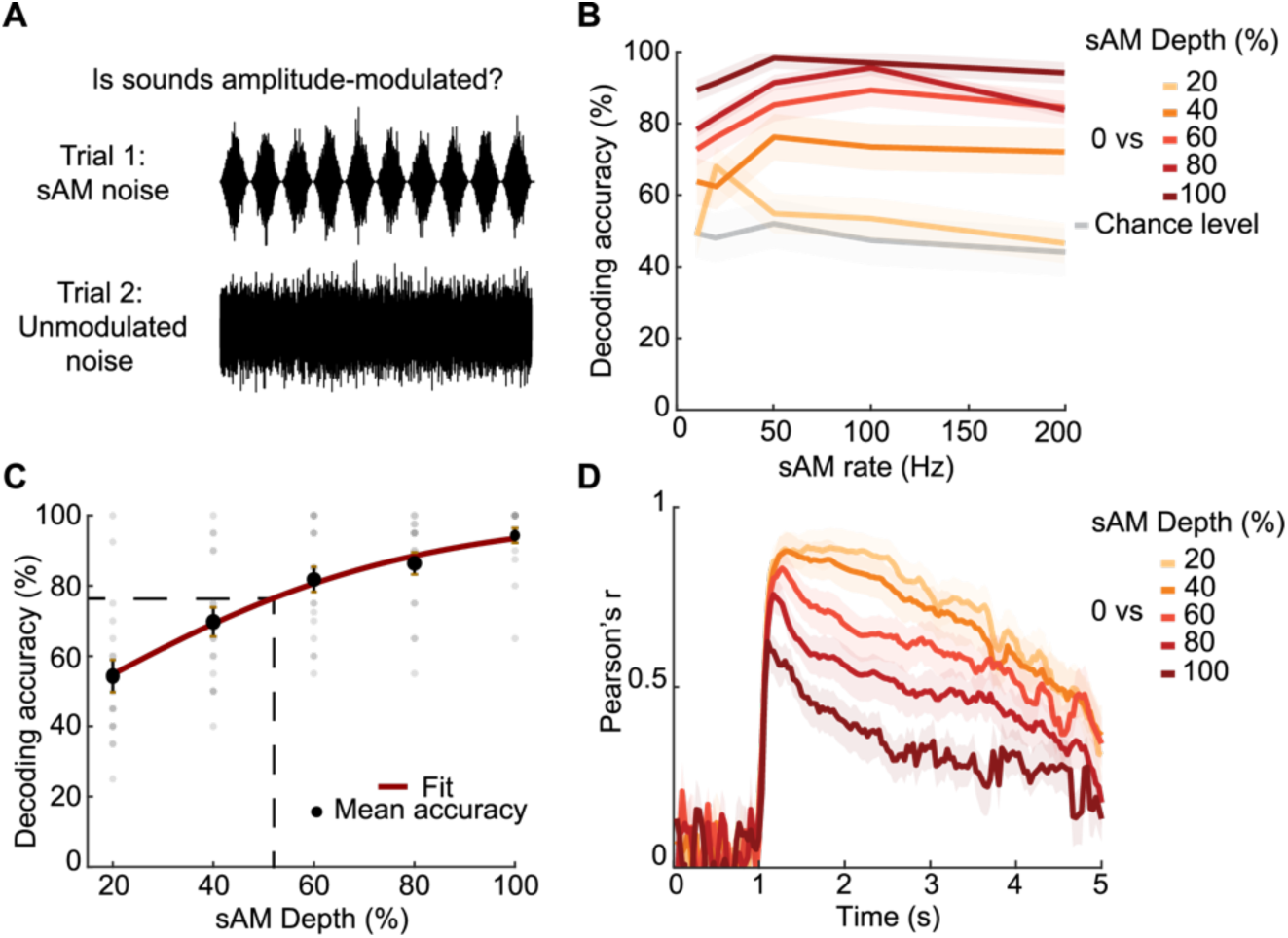
Binary classification of sAM and unmodulated sounds. **A**. Binary classification paradigm: A CNN decoder was trained to classify sAM (with different sAM depths and -rates) and unmodulated noise. **B**. Decoding accuracy of binary classification of sAM sounds and unmodulated noise and corresponding chance level. **C**. Logistic curve fitting of the binary classification performance. Dashed lines, both horizontal and vertical, denotes the position of the half maximum on the fitted curve. **D**. Pattern correlation between sAM sounds with different sAM depths and unmodulated sounds. For **B** and **D**, data are mean ± SEM.

### Minimum resolution of sAM rate representation

Humans, non-human primates, and budgerigars can discriminate 2-5 % rate differences in fully modulated sAM sounds, with rodents showing slightly higher difference limens (Table 1). Given our data showing that fully modulated sAM sounds can be decoded with high accuracy from shell IC neural population activity, we next estimated the sAM rate resolution of shell IC neuron populations. We addressed this question by presenting a series of sAM sounds with narrowly spaced sAM rates (100 % sAM depth, logarithmically spaced between 30-150 Hz with 5 % difference between adjacent stimuli) to a separate group of mice expressing GCaMP8s in shell IC neurons (n = 520 sound-responsive neurons in n = 9 non-overlapping FOVs from N = 5 mice). Individual shell IC neurons had diverse modulation transfer functions similar to our experiments in Figures 1-8 in GCaMP6f/s expressing mice (Figure 9A). Sound-inhibited neurons also had lower neurometric d-prime values compared to sound-excited neurons in these experiments, as observed previously using 6f/6s and more widely spaced sAM rates (Figure 9B; 0.35 ± 0.05 and 1.29 ± 0.07, for sound inhibited and excited neurons, respectively). However, mean d-prime values were similar across the 6f/6s and 8s data sets (Main effect of neuron type: F (1, 1064) = 16.64, p < 0.0001. Main effect of dataset: F (1, 1064) = 2.1, p = 0.1476. Interaction: F (1, 1064) = 0.3125, p = 0.5763, two-way ANOVA). Moreover, the spatial arrangement of sAM rates data points in t-SNE map was a continuous sequence, with similar sAM rates clustering nearby (Figure 9C).

**Table 1.** Behavioral threshold for sAM rate discrimination of fully modulated sAM sounds across different species.

| Source | Subject | sAM rate range | Threshold ($\Delta$ sAM rate) | Performance | Task |
| --- | --- | --- | --- | --- | --- |
| Dooling and Searcy (1981) | Parakeet | 40-160 Hz | 5-12 Hz | — | Detection of increment of sAM rate |
| Long and Clark (1984) | Chinchilla | 20-160 Hz | 8-40 Hz | — | Detection of increment of sAM rate |
| Long and Clark (1984) | Human | 20-160 Hz | 2-10 Hz | — | Detection of increment of sAM rate |
| Formby (1985) | Human | 20-160 Hz | 3-11 Hz | — | Detection of increment of sAM rate |
| Moody (1994) | Macaque | 20-160 Hz | 4-10 Hz | — | Detection of increment of sAM rate |
| Kelly et al. (2006) | Rat | 10-100 Hz | 10-45 Hz | — | Detection of increment of sAM rate |
| Kurt and Ehret (2010) | Mice | 20 and 40 Hz | — | d-prime = 1-1.5 | Two-alternative forced-choice sAM rate discrimination |
| Mackey et al., (2022) | Macaque | 20-120 Hz | 5-10 Hz | — | Detection of increment of sAM rate. ΔsAM rate was varied over a range of 1-64 Hz |
| Yao et al., (2023) | Gerbil | 4 and 10 Hz | — | 90% correct | Two-alternative forced-choice sAM rate discrimination |

Related to Figure 9.

**Figure 9.**
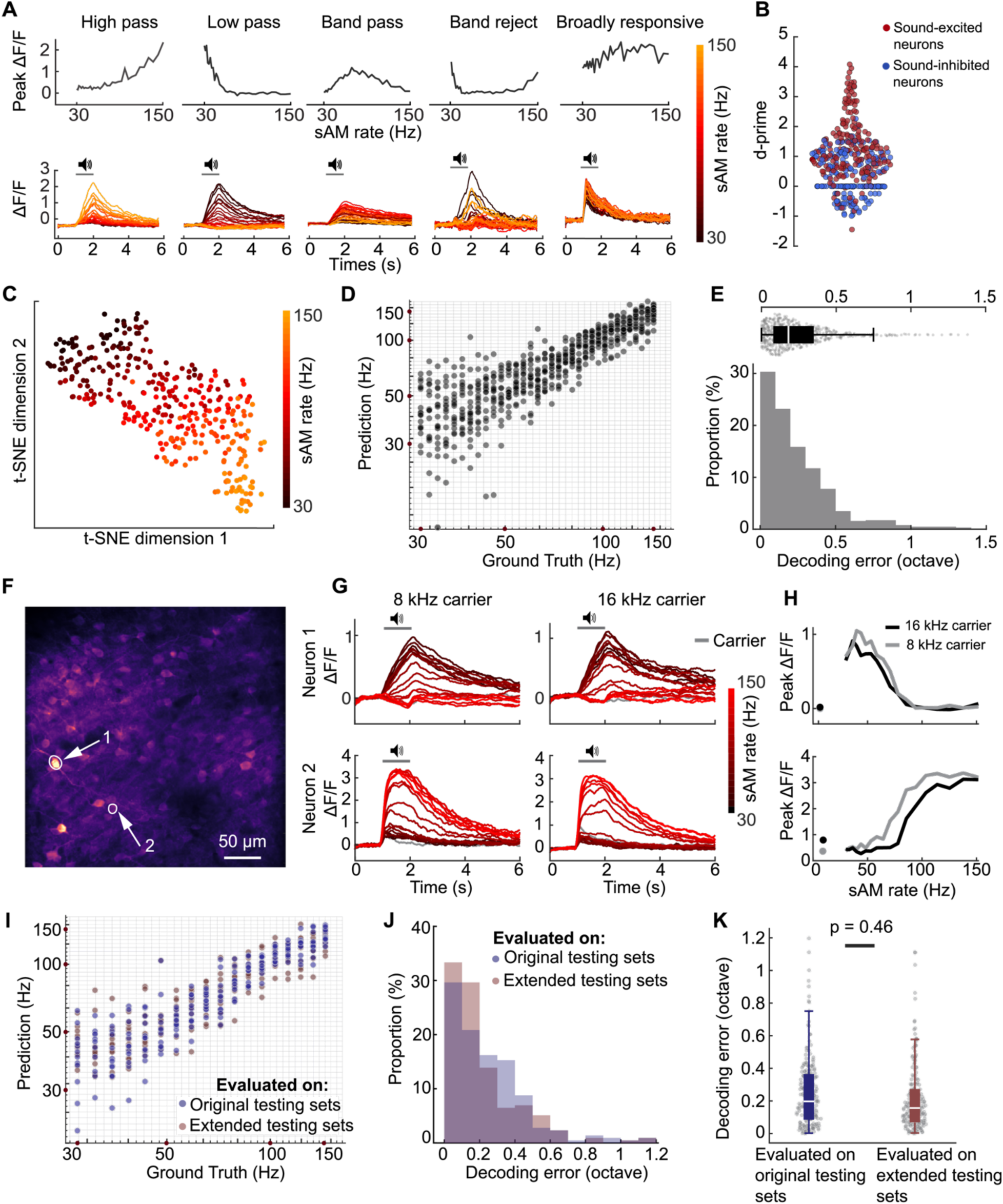
Neuronal responses to narrowly spaced sAM rates in the shell IC. **A**. Examples of modulation transfer functions to fully modulated sAM sounds with 30-150 Hz sAM rates. Top: peak response of example neurons at varying sAM rates and 100% sAM depth. Bottom: trial-averaged fluorescence traces of example neurons. **B**. Distribution of d-prime of shell IC neurons. **C**. t-SNE map of shell IC population responses to narrowly spaced sAM rates. Each dot represents neural population activity in a single trial. **D**. Regression performance of CNN decoder using population fluorescence data of shell IC neurons with narrowly spaced sAM rates and 100% sAM depth. Each dot is the prediction of a single trial in testing sets. The coordinates are plotted on a logarithmic scale of base 10. **E**. Distribution of decoding errors in octaves. **F**. Example imaging FOV. **G-H**. trial-averaged responses (**G**) and peak of response (**H**) of two example neurons to sAM sounds with 16kHz and 8kHZ center frequencies for the noise carrier. **I**. sAM rate decoding performance: sAM rate decoder was trained using data from sAM sounds with central frequency of either 8 kHz or 16 kHz for the noise carrier. After training, the decoder was evaluated on both original held-back testing sets with the same center frequency as in the training set, and extended testing sets with a different central frequency of the carrier. **J**-**K**. Distribution of decoding error in octave. Wilcoxon signed-rank test.

To estimate the lower bound of sAM rate differences that can be transmitted via population codes, we adjusted the CNN decoder to perform a regression task while restricting the architecture to a single convolutional layer. In this paradigm, the median decoding error was 0.18 ± 0.01 octaves (9.12 ± 0.29 Hz; Figure 9D, E). Although few data exist regarding mouse sAM rate discrimination thresholds, the sAM rate resolution in our results is comparable to that of species more sensitive to sound’s temporal characteristics, such as primates and budgerigars (Table 1). We thus expect that sAM rate resolution of shell IC neural population could approximate and may even exceed mouse discrimination thresholds for sAM rates, but further studies are required to directly test this hypothesis.

Finally, we tested if neuronal responses to sAM rates in the shell IC population depend on carrier parameters. However, sAM tuning in single neurons appeared minimally affected when testing two distinct carrier parameters (8 or 16 kHz central frequency of band-limited noise; Figure 9F-H). To further test this idea, we trained the CNN on neural population responses from sAM sounds with either of the two noise carriers. After training, the decoder was tested on either held-out data from the same carrier parameters as data in the training sets, or on extended sets of a different carrier frequency as the training sets. The median decoding error was similar when evaluated on held-out and extended data sets (Figure 9I-K; 0.16 ± 0.01 and 0.20 ± 0.01 octaves, respectively; W = 11031, p = 0.455, Wilcoxon signed-rank test), confirming that the sAM rate representations are minimally impacted by the carrier’s spectral content.

## Discussion

Using *in vivo* 2-photon Ca^2+^ imaging, we determined how sAM rate- and depth are encoded in the activity of shell IC neuron populations. An important consideration is that sAM features can be encoded by average firing rates, temporal synchrony of spikes relative to the sound’s envelope, or a mix of both (Joris et al., 2004). While the slow kinetics of Ca^2+^ signals cannot resolve phase locking of spikes to the sound envelope, peak GCaMP fluorescence is a reasonably good proxy for spike rates across a particular time period (Chen et al., 2013; Ranganathan et al., 2018; Beaulieu-Laroche et al., 2019; Wong and Borst, 2019). Consequently, our data strictly measure the spike rate selectivity of shell IC neurons to sAM features, and we cannot rule out that additional information might be encoded in the temporal patterns of shell IC neuron spiking. When considering this caveat, our decoding results should be considered lower bounds for representational fidelity of sAM features, as any additional temporal information would presumably enhance sAM representations. However, the kinetics of colliculo-thalamic synaptic potentials (∼50 ms halfwidth; Bartlett and Smith, 1999), as well as the membrane time constant of thalamic relay neurons (∼30 ms; Venkataraman and Bartlett, 2013) will likely limit the extent to which any temporal information is transmitted to downstream targets. These collective results further highlight the IC’s role as an important early site for rate coding of time-varying sounds (Dicke et al., 2007).

Individual shell IC neurons were generally broadly tuned to sAM sounds, leading to perhaps unreliable representation of sAM features at the single neuron level. Nevertheless, instead of relying on single neuron activity, information was conveyed more accurately by population responses, in agreement with a recent study showing that ensemble neural discrimination of vocal signals correlates poorly with single unit selectivity in the avian auditory cortex (Robotka et al., 2023). Indeed, sAM rate representational fidelity remained significantly above chance even after excluding individual highly tuned neurons from the neural population, suggesting that the discriminative capacity of in individual neurons is not necessarily a robust index of population-level representations in the shell IC layers. Accordingly, population codes are commonly observed in sensory and motor systems. For instance, mouse piriform cortex represents odor identity based on neural population (Lurilli and Datta, 2017) and population activity of V4 of macaque monkey codes the shape of visual object (Pasupathy and Connor, 2002). Of note, neural population codes are hypothesized to resolve highly precise information sufficient to direct behavior (Partridge et al., 1981; Georgopoulos et al., 1986; Lee et al., 1988; Safaai et al., 2013). Consequently, populations of broadly responsive neurons could effectively transmit multi-dimensional variables that characterize other complex sounds (Zhang and Sejnowski, 1999), such as the fundamental frequency of vowel-like sounds (Carney et al., 2015; Carney, 2018), frequency modulated sweeps, or harmonic stacks.

The fidelity of sAM rate representations depended on sAM depth, with multiple analytical approaches converging upon qualitatively similar estimates of “discrimination threshold” for the shell IC population activity. Specifically, data points in t-SNE mappings of neural population responses to less than 40% sAM depth were barely separable, whereas distinct clusters of sAM rates emerged in the 60% sAM depth mapping plot (Figure 7E). This divergence was mirrored in the pattern correlation analysis across different sAM depths, where there was a sudden drop in pattern correlation between 100% and 40% sAM depths (Figure 7C). sAM rate classifier confusion matrices also agreed with these observations: 60% and 40% sAM depth were in the “watershed area” of CNNs’ decoding performance for sAM rate classification (Figure 4), with a similar estimated sAM *detection* threshold from a binary classification task (∼ 52 % sAM depth; Figure 8). Although future studies will be required to directly test the extent to which shell IC neurons causally contribute to sAM detection or discrimination, the sAM depth thresholds estimates from our neurophysiological data are in qualitative agreement with recently reported behavioral detection thresholds in mice (20-30 % sAM depth; van den Berg et al., 2023). Intriguingly, van den Berg et al. (2023) observed that a higher depth is required for mice to detect sAM sounds as its rate decreases, and this dependence on sAM rate is also mirrored in our binary classification results.

Non-lemniscal thalamic targets of shell IC neurons integrate heterogeneous signals (Lesicko et al., 2020; Liu et al., 2023; Ibrahim et al., 2023), and disynaptically relay this information to limbic circuits that orchestrate learned and innate behaviors (Miura et al., 2020; Carcea et al., 2021; Valtcheva et al., 2023). In contrast to a temporal code, the rate-based encoding of time-varying sounds might allow for a more efficient integration with multisensory, motor, and somatosensory signals across a specific temporal window. Importantly, the temporal characteristics of the sound envelope conveys information to discriminate conspecific vocalizations, and many second-order targets of shell IC neurons are strongly selective for species-specific vocalizations (Gadziola et al., 2012; Grimsley et al., 2013; Gadziola et al., 2016, Hamilton et al., 2021). However, this selectivity may not reflect computations inherent to limbic circuits, but rather could arise upstream in the midbrain: Aitkin et al., (1994) showed that 75 % of feline shell IC neurons respond stronger to vocal stimuli than noise or characteristic frequency stimuli while only ∼25 % of central nucleus neurons do. In mice, many neurons in the low frequency (presumably dorsal) IC regions respond strongly to vocalizations (Portfors et al., 2009). AM features significantly shape the vocalization selectivity in some of these neurons (Holmstrom et al., 2010), and prominent selectivity to ethologically relevant signals is also observed in the shell IC of bats, rats and gerbils (Holmstrom et al., 2007; Gao et al., 2015; Lawlor et al., 2023). In tandem with our current results, these data imply that a shell IC neural population play important roles in processing and integrating vocal signals, thereby shedding light on the perceptual building blocks of behavioral responses in vocal communication.

## Acknowledgments

Funding was provided by the Whitehall Foundation, Hearing Health Foundation, and NIH/NIDCD R01DC019090 to PFA and Magnificent Michigan Undergraduate Research Fellowship to KS. The authors thank Dr. Anahita Mehta for helpful comments.

## Author Contributions

KS, GLQ, MMR, and PFA designed the research. KS, GLQ, ANF, JEC, and PFA conducted the research. KS, GLQ, and MMR analyzed the data. KS, GLQ and PFA wrote the paper.

